# Parallel and sequential pathways of molecular recognition of a tandem-repeat protein and its intrinsically disordered binding partner

**DOI:** 10.1101/2021.04.15.439979

**Authors:** Ben M. Smith, Pamela J. E. Rowling, Chris M. Dobson, Laura S. Itzhaki

## Abstract

The Wnt signalling pathway plays an important role in cell proliferation, differentiation and fate decisions in embryonic development and in the maintenance of adult tissues, and the twelve Armadillo (ARM) repeat-containing protein β-catenin acts as the signal transducer in this pathway. Here we investigate the interaction between β-catenin and the intrinsically disordered transcription factor TCF7L2, comprising a very long nanomolar-affinity interface of approximately 4800 Å^2^ that spans ten of the twelve ARM repeats of β-catenin. First, a fluorescence reporter system for the interaction was engineered and used to determine the kinetic rate constants for the association and dissociation. The association kinetics of TCF7L2 and β-catenin was monophasic and rapid (7.3 ± 0.1 ×10^7^ M^-1^s^-1^), whereas dissociation was biphasic and slow (5.7 ± 0.4 ×10^−4^ s^-1^, 15.2 ± 2.8 ×10^−4^ s^-1^). This reporter system was then combined with site-directed mutagenesis to investigate the striking variability in the conformation adopted by TCF7L2 in the three different crystal structures of the TCF7L2-β-catenin complex. We found that mutation of the N- and C-terminal subdomains of TCF7L2 that adopt relatively fixed conformations in the crystal structures has a large effect on the dissociation kinetics, whereas mutation of the labile sub-domain connecting them has negligible effect. These results point to a two-site avidity mechanism of binding with the linker region forming a “fuzzy” complex involving transient contacts that are not site-specific. Strikingly, two mutations in the N-terminal subdomain that have the largest effects on the dissociation kinetics showed two additional phases, indicating partial flux through an alternative dissociation pathway that is inaccessible to the wild type. The results presented here provide insights into the kinetics of molecular recognition of a long intrinsically disordered region with an elongated repeat-protein surface, a process found to involve parallel routes with sequential steps in each.

## Introduction

β-catenin fulfils two important functions *in vivo*, the first of which is as the signal transducer in the canonical Wnt signalling pathway (Van Der Wal and Van Amerongen, 2020). Upon Wnt stimulation, β-catenin translocates to the nucleus where it binds the T-cell factor/lymphoid enhancer-binding factor (TCF/LEF) family of transcription factors (Arce et al., 2006), and associates with a number of other proteins to form the Wnt enhancesome (Gammons and Bienz, 2018) leading to transcription initiation and elongation and histone and chromatin modification (Mosimann et al., 2009; Willert and Jones, 2006) resulting in the transcription of genes that are of developmental importance and those involved in tissue homeostasis. In the absence of a Wnt signal, cytosolic β-catenin is continuously synthesised then sequestered and targeted for proteasomal degradation by a multi-protein complex called the β-catenin destruction complex (BDC) that forms biological condensates (Schaefer et al., 2018). The BDC comprises five different proteins: two structural proteins Axin and adenomatous polyposis coli (APC), two kinases (glycogen synthase kinase 3β (GSK3β) and casein kinase 1α (CK1α)) and protein phosphatase 2A (PP2A). In the BDC β-catenin is hyperphosphorylated at its N-terminal disordered region by the combined action of GSK3β and CK1α (Kimelman and Xu 2006), allowing it to be recognised and ubiquitinated by the E3 ubiquitin ligase β-TrCP (Wu et al., 2003) and subsequently degraded by the proteasome. The second function of β-catenin is as an adaptor that mediates cell-cell adhesion at adherens junctions, where β-catenin binds to the intracellular domain of the cadherin family of proteins (Harris and Tepass, 2010; Huber and Weis, 2001; Pokutta and Weis, 2007; Van Der Wal and Van Amerongen, 2020). Deregulation of Wnt signalling has been implicated in numerous types of cancer (Bugter et al., 2021) including colorectal and hepatocellular cancers. The deregulation of this pathway results in an increase in transcription which leads to stemness and proliferation. Additionally, disruption of β-catenin binding at adherens junctions is also cancer promoting (Yoshida et al., 2001). Aside from being inherently pro-metastatic through the weakening of cell-cell adhesion (Conacci-Sorrell et al., 2002), adherens junction-associated β-catenin constitutes a significant cellular pool of β-catenin (Herzig et al., 2007), the release of which can result in its nuclear accumulation and unregulated gene activation (Polakis, 2007).

β-catenin has a ∼530-residue central domain of 12 armadillo (ARM) tandem repeats flanked at each end by ∼100-residue disordered regions (Huber et al., 1997; Xing et al., 2008). The twelve imperfect ARM repeats stack linearly to form a right-handed superhelix of helices. The third helix of each ARM repeat lines the groove formed by the superhelix and is enriched in positively charged residues (Huber et al., 1997), creating an extended docking site for β-catenin’s negatively charged binding intrinsically disordered partners APC, E-cadherin and the TCF/LEF family of transcription factors that wrap around it (Eklof Spink et al., 2001; Graham et al., 2000; Ha et al., 2004a; Poy et al., 2001; Xing et al., 2004) (Fig. 1).

**Figure 1.**
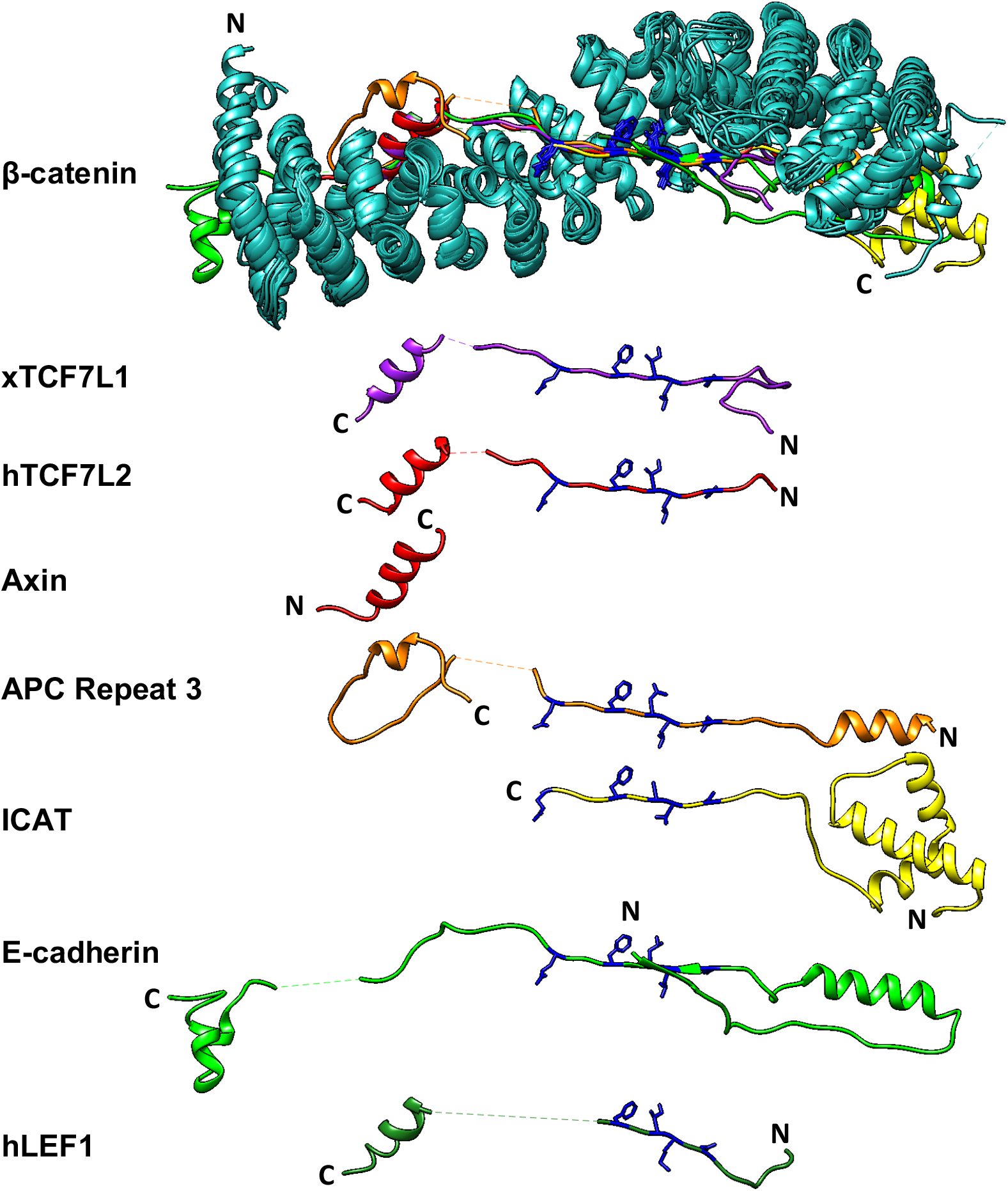
Interactions of β-catenin and its partner proteins: Schematic showing the superposition of the crystal structures of β-catenin (cyan) in complex with its seven binding partners (various colours), visualised in UCSF Chimera: *Xenopus* TCF7L1 (PDB ID 1G3J, Graham *et al*. (2000)), human Tcf4 (2GL7, Sampietro *et al*. (2006)), Axin (1QZ7, Xing *et al*., (2003)), APC third repeat (1TH1, Xing *et al*. (2004)), ICAT (1LUJ, Graham *et al*. (2002)), the cytoplasmic domain of E-cadherin (3IFQ, Choi *et al* (2009)), and human Lef-1 (3OUW, Sun and Weis (2011)). Disordered regions that are not visible in the crystal structures are denoted by a dashed line. All partner proteins, except for Axin, bind in an antiparallel configuration relative to β-catenin. The consensus binding motif DxΦΦxΩx_2-7_E, is highlighted (dark blue), where Φ indicates a hydrophobic residue and Ω ian aromatic residue.

The target genes for the canonical Wnt pathway have a conserved sequence that binds to members of the TCF/LEF family of transcription factors, of which Transcription factor 7-like 2 (TCF7L2, also known as Tcf4) has been the most widely studied to date (Cadigan and Waterman, 2012; Schuijers et al., 2014). TCF7L2 mutations are implicated in many types of cancers (Tang et al., 2008) and have been shown to promote migration and invasion of human colorectal cancer cells (Hazra et al., 2008; Slattery et al., 2008; Wenzel et al., 2020). Single nucleotide polymorphisms within the TCF7L2 gene are also a genetic biomarker for an increased risk of both type 2 and gestational diabetes (Vaquero et al., 2012; Zhang et al., 2013). TCF7L2 is like many mammalian transcription in having an intrinsically disordered region (IDR) The absence of fixed tertiary structure is thought to give proteins conformational plasticity and enable them to bind to a number of different macromolecules in response to physiological needs (Borgia et al., 2018; Csizmok et al., 2016; Holmstrom et al., 2019; Van Der Lee et al., 2014; Schuler et al., 2019; Shammas, 2017; Tsafou et al., 2018). Given their capacity to adjust to any given binding partner, the range of structures adopted by IDRs upon complex formation is unsurprising. Broadly, these can be divided into static complexes (Uversky, 2011) where the IDR is ordered (and visible in X-ray crystal structures), and dynamic or “fuzzy” complexes, where the IDR retains a significant proportion (sometimes all) of its disorder in the complex (Borg et al., 2007; Mittag et al., 2009; Sigalov et al., 2007). IDP binding mechanisms are equally varied, and they have grouped them into four distinct classes: simple two-state binding, avidity, allovalency, and fuzzy binding (Olsen et al., 2017). Even in the more simple case of two-state binding, association can involve conformational selection or folding upon binding (Bachmann et al., 2011; Dogan et al., 2014; Karlsson et al., 2012; Rogers et al., 2014; Wright and Dyson, 2009).

Intriguingly, there are three crystal structures of TCF7L2 in complex with the ARM repeat domain of β-catenin, with TCF7L2 adopting a different conformation and making different β-catenin contacts in each (Fig. 2) (Graham et al., 2000, 2001; Poy et al., 2001; Sampietro et al., 2006). In addition, there are significant portions of missing residues and different regions are resolved in the different structures, suggesting that TCF7L2 is highly dynamic in complex with β-catenin. Similar behaviour is observed in the crystal structures of other β-catenin complexes, providing tantalising hints as to the underlying mechanisms of recognition (Choi et al., 2009; Daniels and Weis, 2002; Graham et al., 2002; Ha et al., 2004; Huber and Weis, 2001; Xing et al., 2003, 2004). Most recently, a single-molecule study by Hofmann and co-workers of the interaction between β-catenin and E-cadherin revealed a rugged energy landscape of E-cadherin with many shallow minima, a fuzzy complex, and a mechanism of intrachain diffusion of E-cadherin on β-catenin (Wiggers et al., 2021). Here, we use site-directed mutagenesis and kinetic analysis to explore the significance of this plasticity in the β-catenin-TCF7L2 complex and define the mechanism of association between the two proteins.

**Figure 2.**
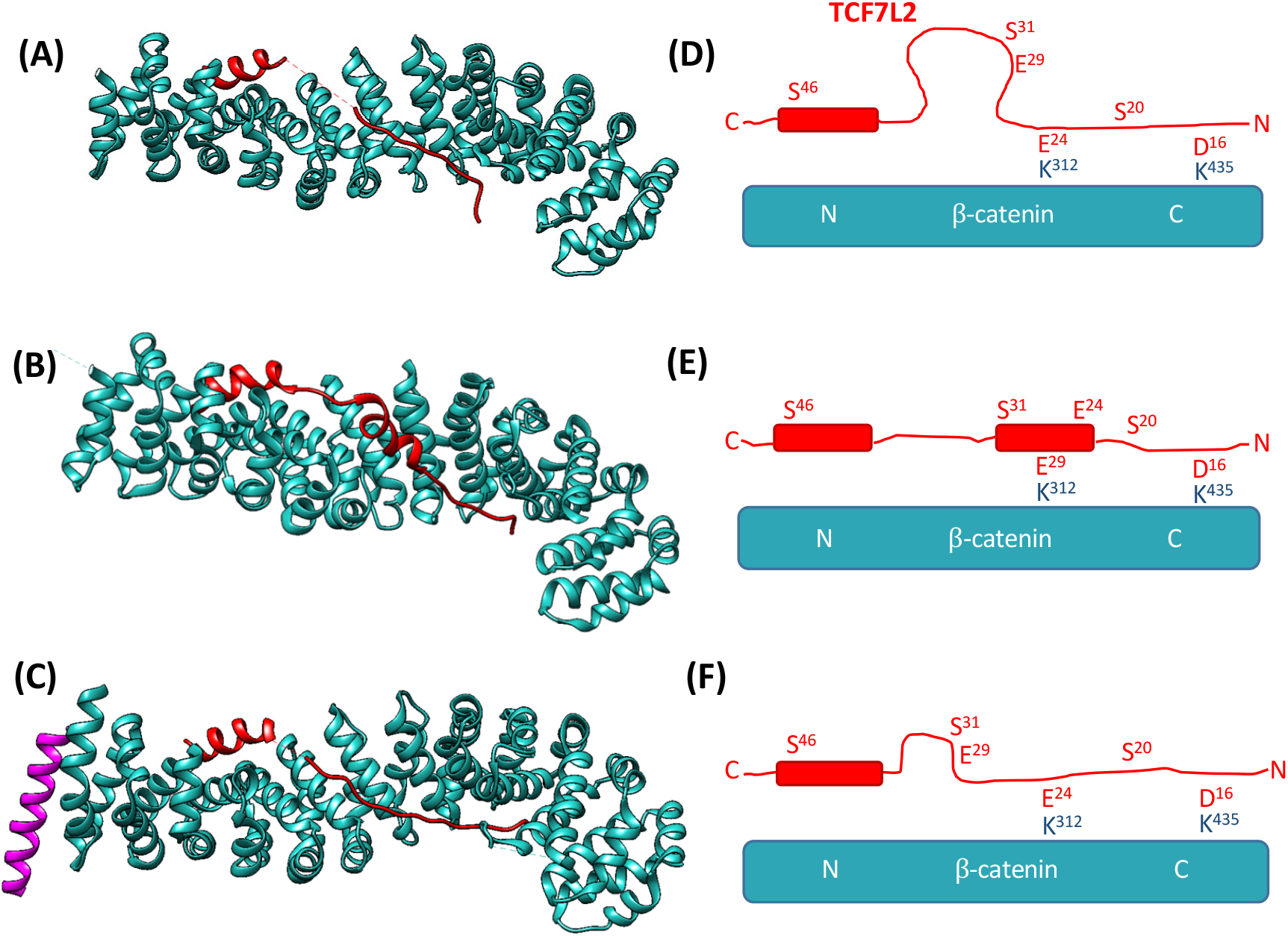
Schematics showing the crystal structures of the β-catenin-TCF7L2 complex. (A), (B) and (C) show the structures of TCF7L2 (red) in complex with β-catenin (cyan) obtained by Poy *et al*., Graham *et al*. and Sampietro *et al*. respectively. Structure (C) also has a peptide from the protein BCL9 (purple) bound. (D), (E) and (F) are schematics highlighting the different conformations of TCF7L2 between the three structures and indicating the positioning of the key contact residues within each structure. Images (A) (B) and (C) were created using UCSF Chimera (Pettersen et al., 2004) from PDB 1JPW (Poy et al., 2001), 1JDH (Graham et al., 2001) and 2GL7 (Sampietro et al., 2006), respectively.

## Materials and Methods

### Molecular biology and protein expression and purification

Mutations in the β-catenin binding fragment of TCF7L2 (1-54 amino acids) were introduced by site directed mutagenesis. The plasmid encoding the armadillo (ARM) repeat domain of human β-catenin (residues 134-671) was a kind gift from Prof. W.I. Weis. All proteins were expressed in *E. coli* C41 cells (Miroux and Walker, 1996). Transformed cells were grown in 2TY with appropriate antibiotic until OD600 of 0.6 at 37 °C, the temperature was then lowered to 25 °C and the cells induced with 0.2 mM IPTG after a further 18 h the cells were harvested at 5000 g for 7 min at 4 °C. The cells were resuspended in 50 mM Tris pH7.5, 150 mM NaCl, 1 mM DTT containing protease inhibitors and lysed using an Emusiflex C5 (Avestin) at 10 000 psi. The lysed cells were centrifuged at 35 000 g for 35 min at 4 °C.

The GST-tagged β-catenin supernatant was incubated with glutathione-Sepharose 4B (Cytiva) and washed to remove unbound proteins. The GST tag was cleaved from the target protein on resin using thrombin and the protein eluted. β-catenin was further purified using a Mono-Q column (GE Life Sciences) equilibrated in 50 mM Tris-HCl buffer pH 8.9, 50 mM NaCl, 1 mM DTT and eluted with a linear NaCl gradient to 1 M NaCl. Fractions containing greater than 95% β-catenin were pooled snap frozen and lyophilised and stored at -80 °C.

The His-tagged TCF7L2 constructs were affinity purified using Ni-NTA agarose (Qiagen) and the tagged cleaved from the protein using thrombin. THE TCF7L2 was further purified on a HiLoad 26/600 Superdex 75 pg (Cytiva) equilibrated in PBS containing 1 mM DTT. The fractions corresponding to A_215_ peaks were analysed by SDS-PAGE and the fractions containing TCF7L2 at greater than >95% were flash frozen with liquid nitrogen and stored at -80 °C until use.

The identities of all proteins were confirmed by MALDI Mass Spectrometry performed by Dr. Len Packman (University of Cambridge PNAC Facility). Protein concentration of β-catenin was calculated from its extinction coefficient obtained using ProtParam (Gasteiger et al., 2005) at 280nm and TCF7L2 proteins concentration was measured using the Pierce™ BCA protein Assay Kit (ThermoFisher) ion-exchange-ninhydrin analysis performed by Dr. Peter Sharratt (University of Cambridge PNAC Facility).

### Fluorescent labelling of TCF7L2

The maleimide functional group reacts with sulfhydryl groups via nucleophilic conjugation addition and was used to label proteins at cysteine residues. A 500 µl aliquot of TCF7L2 (1-54) construct containing a cysteine residue buffer exchanged into PBS. A stock solution of 20 mM fluorescein maleimide in 100% DMSO was added at a 5-times molar excess and incubated at 37°C in the dark for hr. Excess fluorescein maleimide was removed by the method of Vivès and Lebleu, (2003). The final precipitated protein pellet was left at room temperature, and open to the air for 16 hrs to ensure that any residual acetone had evaporated, after which the desiccated labelled-TCF7L2 was stored at 4 °C for up to 5 days without loss of activity.

### Isothermal titration calorimetry (ITC)

ITC measurements were performed using the VP-ITC calorimeter (Malvern Pananalytical). Both β-catenin and TCF7L2 constructs were stored as lyophilised powders after labelling. To ensure matched buffers, lyophilised β-catenin was resuspended in ultra-pure water and the buffer exchanged into 50 mM Tris-HCl buffer pH 8.9, 100 mM NaCl, 2 mM DTT and the TCF7L2 was resuspended in the same buffer. Unlabelled TCF7L2 was washed with acetone three times, as described previously to produce a lyophilised pellet. β-catenin was used at a concentration of 2-4 µM in the cell and TCF7L2 was used at 6-10x higher concentration in the syringe (18-40 µM). Blank titrations of TCF7L2 into buffer and buffer into β-catenin were performed to demonstrate that the reaction observed was a binding interaction. All experiments were performed at 30°C. Data were fitted the Origin software package supplied with the instrument using the one-site binding equation.

### Kinetic experiments

Stopped-flow spectroscopy was performed using a SX-19 Stopped flow fluorimeter (Applied Photophysics) in fluorescence mode. Excitation and emission wavelengths were set to 495 nm and 519 nm respectively and a 515 nm long-pass filter was used to improve the signal to noise ratio. All proteins were prepared in fresh PBS buffer, 1 mM DTT, and the experiments performed at 15°C. The β-catenin concentrations were at least ten times higher than that of the TCF7L2 to ensure a pseudo-first order association reaction; a fixed concentration of fluorescently labelled TCF7L2 (10 nM) construct was rapidly mixed, in a 1:1 volume ratio, with varying concentrations unlabelled β-catenin (between 100 nM and 1000nM) and the change in fluorescence intensity was recorded. For dissociation experiments, a complex of labelled TCF7L2 and β-catenin was pre-formed by mixing to two proteins in a 1:1 molar ratio (200 nM) and incubating them at 15°C for 1 hr in the dark to reach equilibrium. The pre-formed complex was then mixed in a 1:1 volume ratio with 50 times molar excess unlabelled wild-type TCF7L2 (10 μM), and the change in fluorescence intensity was measured. At each β-catenin concentration a minimum of three traces were collected and averaged. The average was plotted using GraphPad Prism 6 (GraphPad Software, Ltd), and fitted to either a single-exponential phase or a two-phase exponential.

For the association experiments, the observed rate constants, *k*_*obs*_, were plotted against the concentration of β-catenin and *k*_*on*_ calculated from the linear fit:

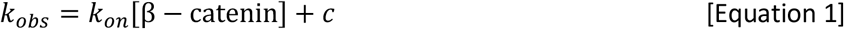

For the slow dissociation kinetics, fluorescence spectroscopy was performed using a LS55 Luminescence Spectrometer (Perkin-Elmer) using a cell of pathlength 10 mm. Excitation and emission wavelengths were set to 495 nm and 519 nm respectively, both excitation and emission shutter width were 5 nm and photomultiplier tube was set to 650 V. All proteins were prepared in fresh 1x PBS buffer with 1 mM DTT and the experiments performed at 15 °C, except when otherwise stated. A complex of labelled TCF7L2 and β-catenin was pre-formed by mixing to two proteins in a 1:1 molar ratio and incubating them at 15 °C for 1 hr in the dark to reach equilibrium. The pre-formed complex was then mixed in a 1:1 ratio with 50 times molar excess unlabelled WT TCF7L2 and the change in fluorescence intensity was measured. Individual traces were fitted as described above.

## Results

### The association of TCF7L2 binding with β-catenin is monophasic, and dissociation of the complex is biphasic

In order to study the kinetics of TCF7L2-β-catenin complex formation and dissociation, we used a recombinant fragment of TCF7L2 comprising the N-terminal 54-residue β-catenin binding domain. We made a conservative serine to cysteine mutation in the wild-type TCF7L2 (1-54) to allow us to label the protein at a unique site. The wild-type TCF7L2 (1-54) construct contains eight serine residues, excluding the one left behind as a result of removing the purification tag with thrombin. The crystal structures were inspected to find a position where the addition of a bulky, hydrophobic dye molecule would not have a significant steric effect on binding but would be able to report on complex formation. Contact map analysis revealed that only one serine residue, S^31^, formed no contacts to β-catenin in any of the crystal structures, indicating that it is in a sterically unhindered position. Consequently, S^31^ was mutated to a cysteine for labelling with fluorescein maleimide at this position. The TCF7L2 (1-54) S31C construct is hereafter referred to as “WT” and is used as a pseudo-wild type with which all other TCF7L2 mutants are compared. In order to determine whether the S31C mutation, and its subsequent labelling with fluorescein maleimide, had a significant effect on TCF7L2 binding to β-catenin, equilibrium binding experiments were performed using ITC. Previous studies have used ELISA (Omer et al., 1999) and ITC (Sun and Weis, 2011) to determine the binding affinity between β-catenin and a similar TCF7L2 fragment. The values they obtained were 15 ± 6 nM and 16 ± 3 nM for TCF7L2 (1-53) and TCF7L2 (1-57), respectively. The results of the ITC experiments showed that there was no significant difference between TCF7L2 (1-54), WT, fluorescein-labelled WT and the literature values (Table S1 and Fig. S1).

Association kinetics were monitored by the change in fluorescence intensity of the conjugated fluorescein moiety in a stopped-flow fluorimeter. Dissociation kinetics were monitored using either a stopped-flow fluorimeter or a fluorescence spectrometer depending on the timescale of the reaction. Association kinetics were initiated by rapid mixing in the stopped-flow of solutions containing a fixed concentration of fluorescein-labelled WT, and varying concentrations of β-catenin, in order to produce a pseudo-first-order regime. All of the association traces showed a mono-exponential decrease in fluorescence with time, indicating a single association event (Fig. 3). Additionally, the observed association rate (*k*_*obs*_) increased linearly with increasing β-catenin concentration confirming that the reaction is bimolecular. The association rate constant, *k*_*on*_, calculated from the pseudo-first-order plot, was 7.33 ± 0.14 ×10^7^ M^-1^ s^-1^.

**Figure 3.**
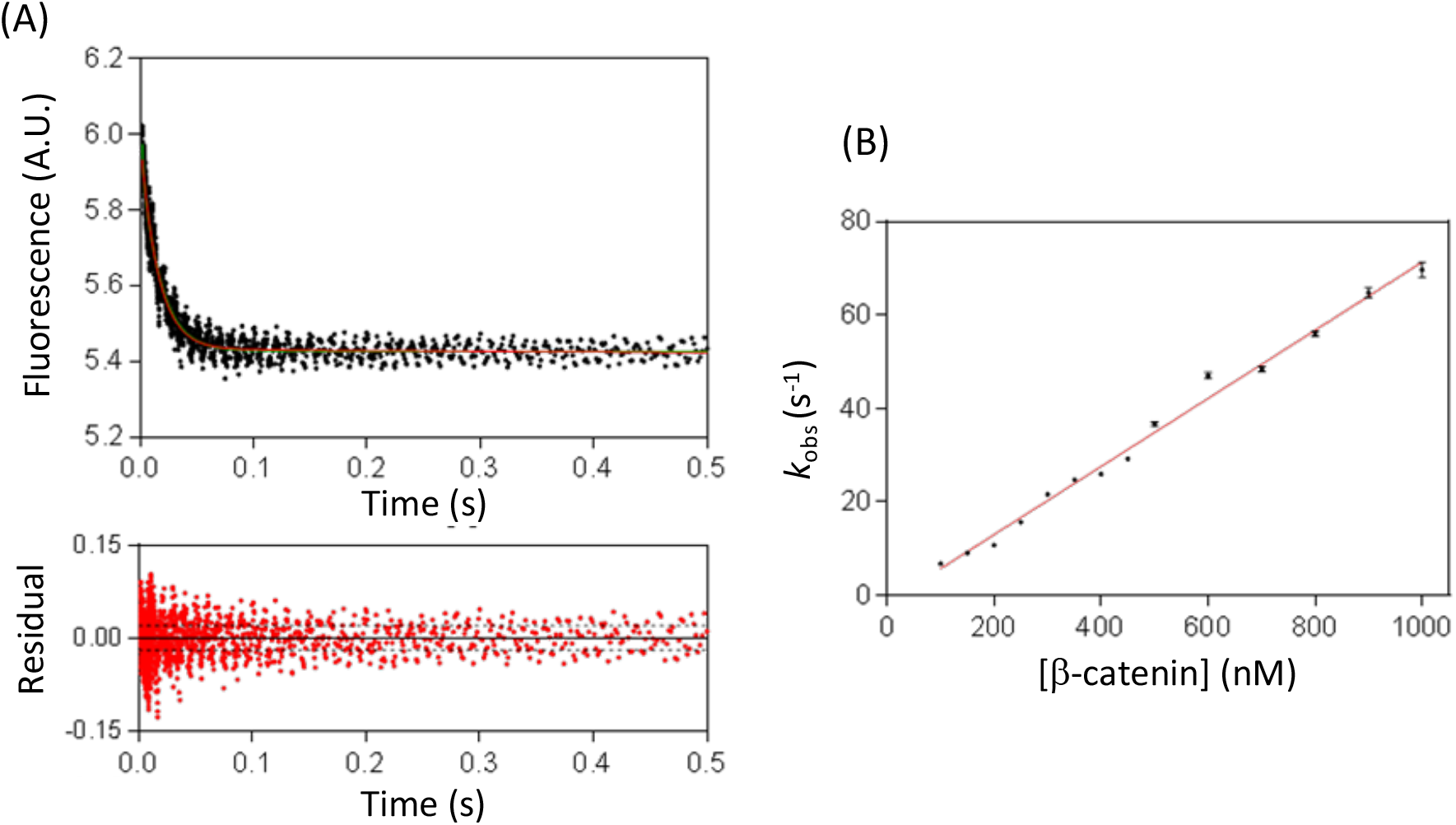
Stopped-flow fluorescence measurement of the association kinetics of TCF7L2 and β-catenin. (A) Time-dependent change in fluorescence upon mixing of 10 nM fluorescently labelled TCF7L2(1-54) and 900 nM β-catenin. The data were fitted to a single-exponential function and the residuals are shown below the main plot. (B) The observed rate constant is plotted as a function of the concentration of β-catenin, from which the association rate constant, *k*_*on*_, is calculated. The errors bars in (B) are the standard deviations of the fits. Experiments were performed in PBS buffer, 1 mM DTT at 15°C.

Dissociation kinetics were monitored by first forming a 1:1 complex of β-catenin and fluorescent-labelled WT-TCF7L2. The complex was dissociated by mixing with excess unlabelled TCF7L2 (1-54). Dissociation traces showed a biphasic increase in fluorescence intensity demonstrating that, unlike association, dissociation is a two-step process (Fig. 4). The traces did not reach a plateau when monitored by stopped-flow (Fig. 4A), and consequently a fluorimeter was used instead (Fig. 4B). The data were fitted to the sum of two exponential phases, comprising a major, slow phase with *k*_*off,major*_ = 5.73 ± 0.40 ×10^−4^ s^-1^ (relative amplitude of 77 ± 3 %) and a minor, fast phase with *k*_*off,minor*_ = 1.52 ± 0.28 ×10^−3^ s^-1^ (relative amplitude of 23 ± 3 %). The dissociation kinetics were independent of β-catenin concentration.

**Figure 4.**
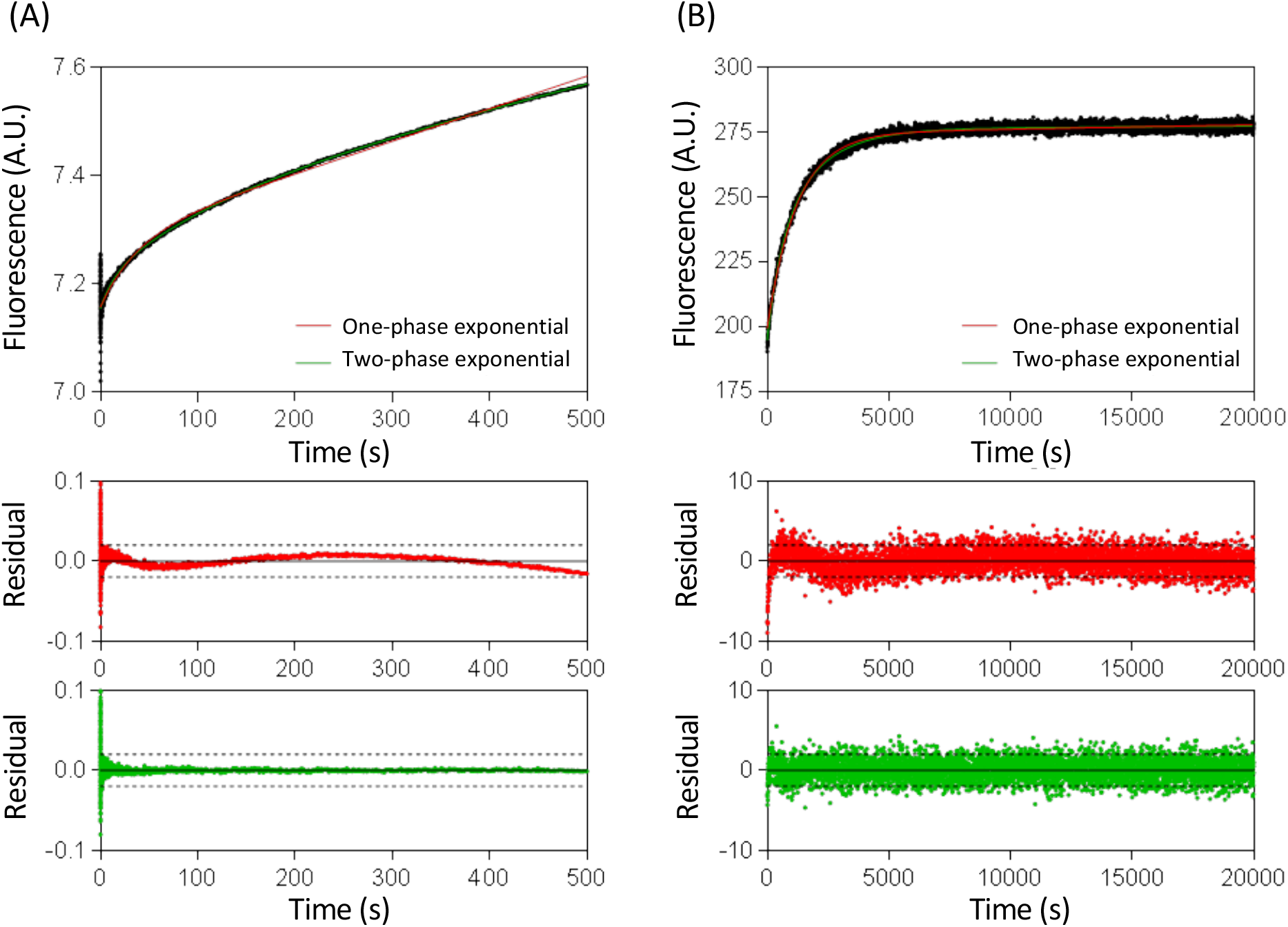
Stopped-flow and time-dependent fluorescence spectroscopy measurements of the dissociation of the WT TCF7L2-β-catenin complex. 200 nM of fluorescent-labelled WT-TCF7L2-β-catenin complex was mixed in with 50 times molar excess of unlabelled WT TCF7L2 (1-54) and the reaction was monitored by (A) stopped-flow fluorescence and (B) fluorescence. The traces were fitted to a single exponential phase and to the sum of two-phase exponential phases, and the corresponding residuals are plotted below. Experiments were performed in PBS buffer, 1 mM DTT at 15°C.

### Alanine scanning of TCF7L2 reveals three distinct interface regions

To investigate the contribution of individual residues to the kinetics of the TCF7L2-β-catenin interaction, an alanine scan of TCF7L2 was performed. Residues spanning the length of the β-catenin binding interface were mutated to alanine. The association traces of all of the mutants showed a single-exponential decrease in fluorescence intensity, similar to that of WT, and none had a significant effect on the association rate constant (Table 1). Several mutants had a very large effect on the dissociation kinetics (Table 1). For I19A, the dissociation kinetics was sufficiently fast as to be measurable on the stopped-flow instrument as well as on the fluorimeter. For I19A, F21A, L41A and V44A, the dissociation kinetics were too fast for the fluorimeter, and the stopped-flow was used instead. For all mutants except I19A and F21A, which are discussed in more detail below, the dissociation traces were biphasic with a major slow phase of 70-90% and a minor fast phase of 10-30%, similar to those of WT.

**Table 1:**
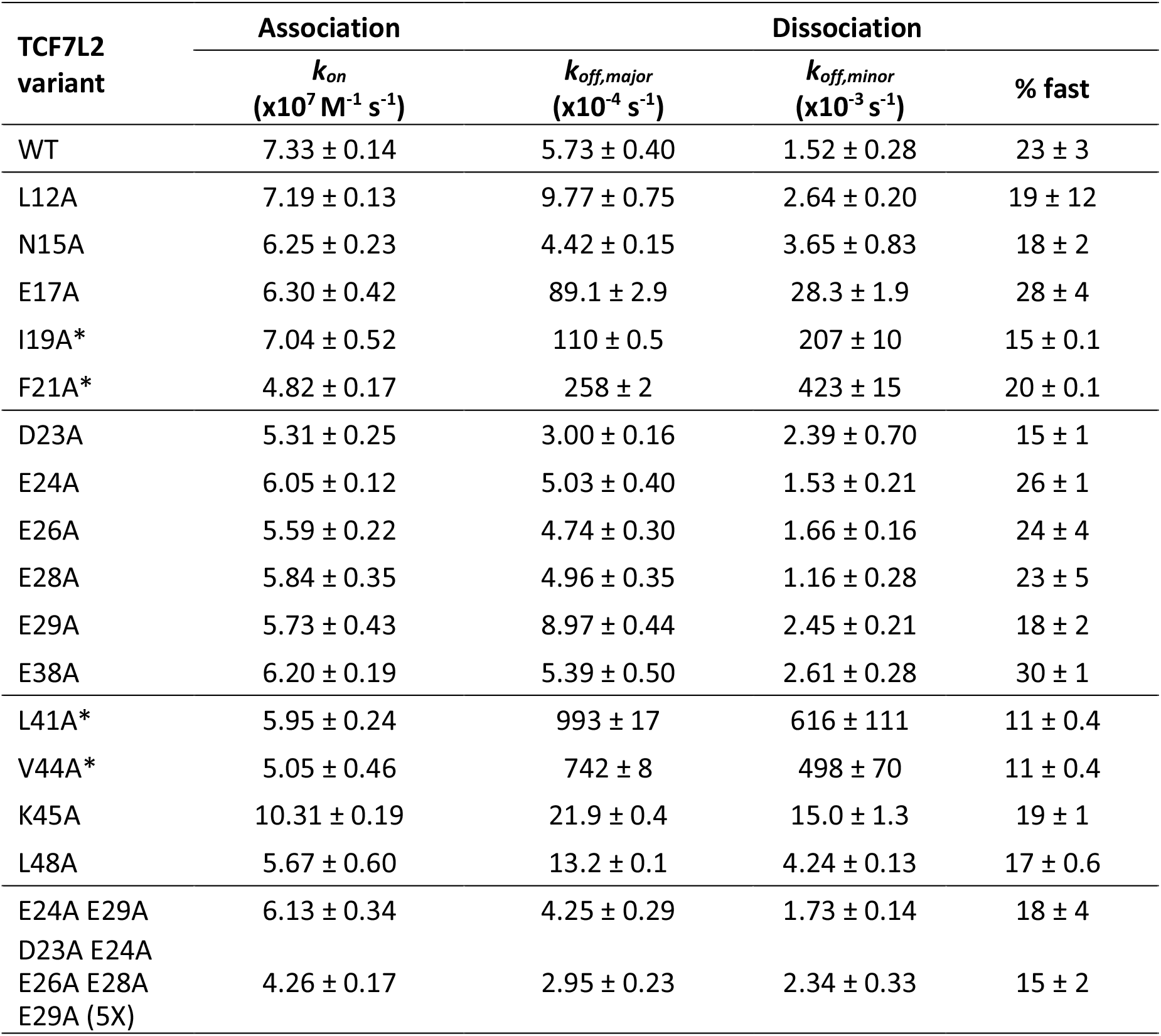
Association and dissociation kinetics for the interaction of alanine mutants of TCF7L2. All mutations were constructed in the S31C variant (denoted WT) and were labelled with fluorescein at this position. Association kinetics were measured on a stopped-flow fluorimeter and dissociation kinetics on a fluorimeter. *The exceptions are I19A, F21A, L41A and V44A, for which the data listed for association and dissociation kinetics were obtained on the stopped-flow.

From these data, we were able to divide TCF7L2 into three distinct regions, which we refer to as the N-terminal binding region (residues 12-22), the variable region (residues 23-39) and the C-terminal binding region (residues 40-50) (Fig. 5). Mutations in the N- and C-terminal binding regions have a much larger effect on the TCF7L2-β-catenin kinetics than do mutations within the variable region, and a similar trend was observed previously for the equilibrium binding (Knapp et al., 2001; Omer et al., 1999). The contributions of the three regions to binding affinity appear to mirror their conformational properties *in crystallo* in that the N- and C-terminal regions, which contribute most to the binding affinity, adopt a relatively ‘fixed’ conformation that is similar in all three crystal structures, whereas the variable region is more flexible or ‘pliable’ and adopts different conformations in the three structures and makes little contribution to binding affinity.

**Figure 5.**
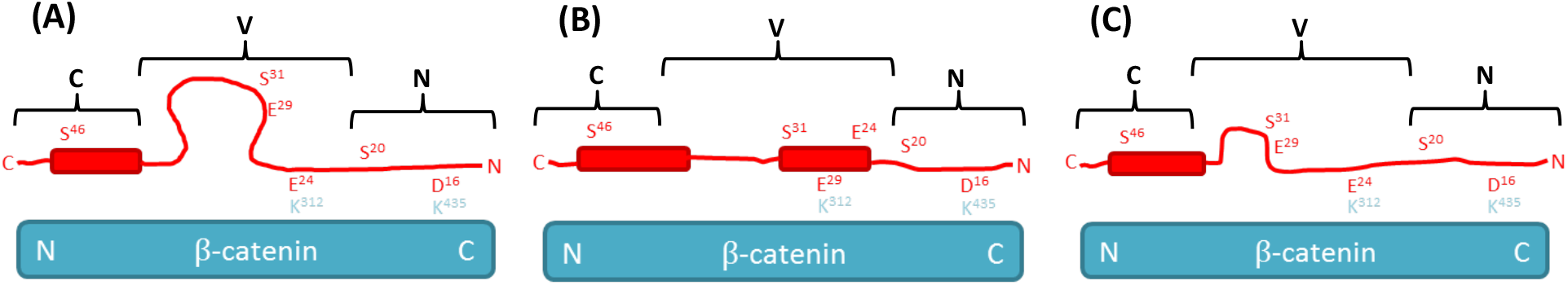
Schematic showing the three interface regions (N, variable (V), and C) of TCF7L2. Schematics showing the conformational preferences of TCF7L2 (red) in complex with β-catenin (cyan) in the different crystal structures. (A), (B) and (C) are based on the crystal structures of Poy *et al*., Graham *et al*. and Sampietro *et al*., respectively. The N-terminal binding domain (N) and the C-terminal binding domain (C) each adopt very similar conformations in all three structures, but the variable region (V) has a different conformation in each structure.

### Multi-site mutations in the variable region of TCF7L2 do not perturb the kinetics

Next, we sought to explore further the role of the variable region. The Sampietro structure is most similar to the Poy structure, and both show Glu24 of TCF7L2 interacting with Lys312 of β-catenin (Fig. 5A,C). In the Graham structure, it is Glu29 that interacts with Lys312; this difference appears to be due to residues 21-32 of TCF7L2 forming an α-helix rather than an extended conformation, forcing Glu24 to point away from the surface of β-catenin and positioning Glu29 in the equivalent position where it can interact with Lys312 (Fig. 5B). Other studies of β-catenin complexes have identified Lys312 and Lys435 as key interaction residues (often referred to as the “lysine buttons” or “charged buttons”), and so it seems strange that there should be such variation in the TCF7L2 structure at this critical interface with β-catenin (Graham *et al*., 2000; Gail, Frank and Wittinghofer, 2005; Xu and Kimelman, 2007; Sun and Weis, 2011). In the Graham structure, the entire variable region of TCF7L2 (23-39) is visible, whereas in the other two structures a significant number of residues are disordered and not visible (residues 27 to 39 in the Poy structure, and residues 29 to 37 in the Sampietro structure). Additionally, the C-terminal α-helix (residues 41-49), which is present in all three structures, has an extra turn in the Graham structure (residues 37-40). These more extensive helical stretches of the Graham structure have the effect of pulling the variable region across the β-catenin surface. We noticed that there are a further three Glutamate residues close to Glu24 and Glu29 that could potentially interact with Lys312 of β-catenin: Glu23, Glu26 and Glu28, and we postulated that conformations may be populated other than those that are observed *in crystallo*. By altering the location and length of the N-terminal α-helix of Graham’s structure, the register would be altered and Glu26 or Glu28 could be presented to the lysine button. To investigate this possibility, two additional alanine variants were made, the double mutant E24A E29A and the quintuple mutant D23A E24A E26A E28A E29A (referred to subsequently 5X). We were surprised to find, given the anticipated key interaction of Glutamate with the lysine button, that neither the double mutant nor the quintuple mutant had a significant effect on the kinetics.

### TCF7L2 mutants I19A and L21A display more complex dissociation kinetics

The dissociation kinetics of the TCF7L2 I19A mutant occurred over a faster timescale than WT and could be measured on both the stopped-flow instrument and the fluorimeter (Fig. 6). On the stopped-flow, the kinetics traces were captured adequately by a biphasic equation with a minor fast phase of 15% of the total amplitude and a major slow phase of 85%, in line with the other mutants. A biphasic equation was also used to fit the kinetic traces obtained on the fluorimeter, with a major fast phase of 82% amplitude and a minor slow phase of 18% amplitude. The slower phase obtained from stopped flow and the faster phase obtained from the fluorimeter have similar rate constants (11.0 ×10^−3^ s^-1^ and 16.5 ×10^−3^ s^-1^ respectively), suggesting that the dissociation of I19A comprises a least three phases in total.

**Figure 6.**
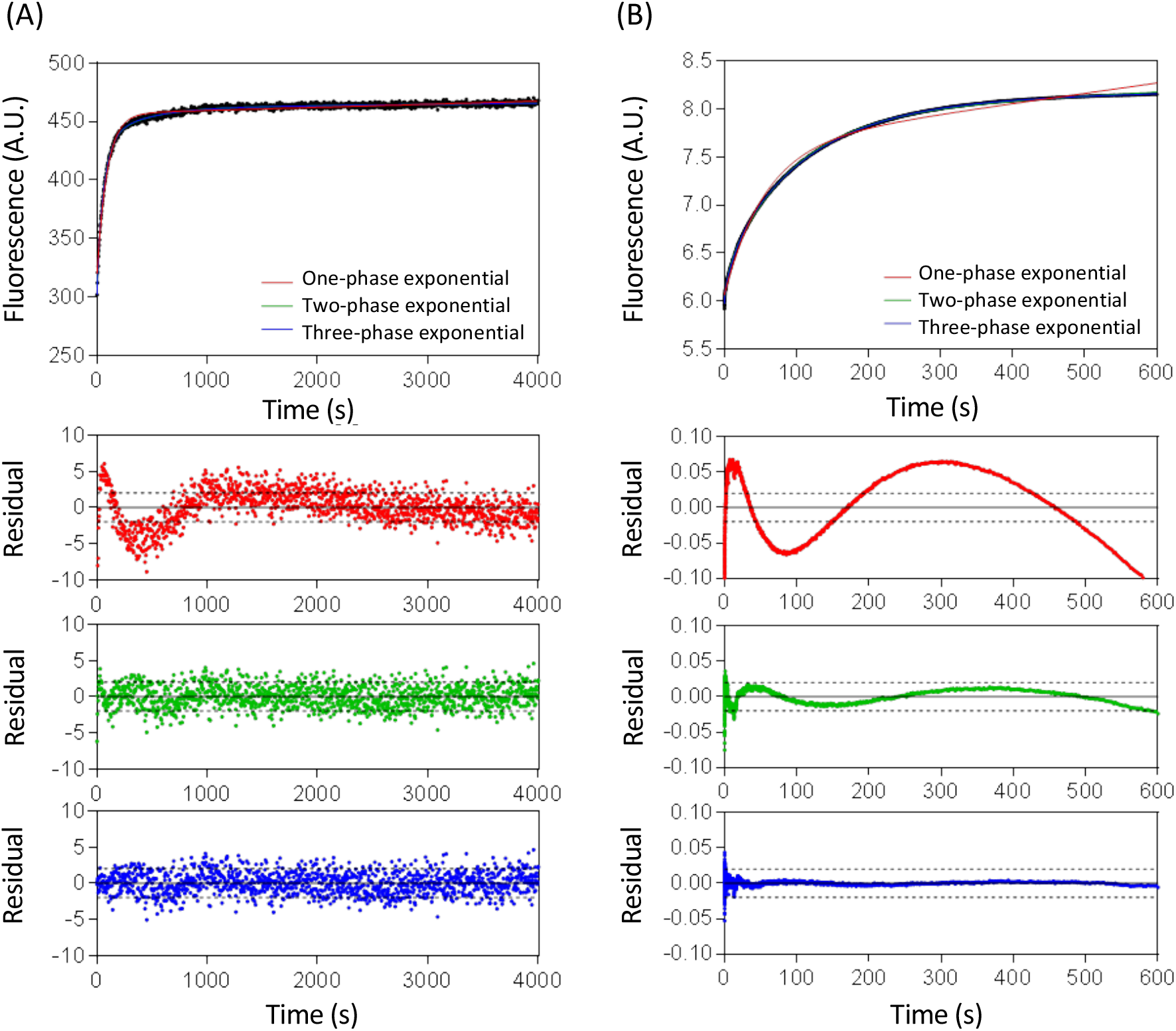
Dissociation kinetics of TCF7L2 I19A. (A) and (B) are representative dissociation traces for I19A obtained using the fluorimeter and the stopped-flow instrument, respectively. The data were fitted to one-, two- and three-phase exponential functions, and the residuals are also shown (red, green and blue respectively). Experiments were performed in PBS buffer, 1 mM DTT at 15°C.

We next attempted to fit the dissociation kinetics of TCF7L2 I19A to a three-phase exponential function, and the rate constants and relative amplitudes obtained are listed in Table 2. Using the fluorimeter, the major phase (75%) had a rate constant of 15.0 ± 0.8 ×10^−3^ s^-1^, and there was a minor (15%) slow phase of 2.7 ± 0.1 ×10^−3^ s^-1^ and a minor (10%) fast phase of 88 ± 19 ×10^−3^ s^-1^. Using the stopped flow, the major phase (79%) was the slowest phase, with a rate constant was 9.1 ± 0.2 ×10^−3^ s^-1^, and there were two faster minor phases with rate constants of 65.2 ± 7.3 ×10^−3^ (15%) and 535 ± 70 ×10^−3^ s^-1^ (16%) and the minor middle phase. These results indicate that there are at least three and likely four phases in the dissociation of I19A from β-catenin.

**Table 2:** Multi-phase dissociation kinetics of TCF7L2 I19A.

| Instrument | $k_1$<br>($\times 10^{-3} \text{ s}^{-1}$) | $A_1$<br>(%) | $k_2$<br>($\times 10^{-3} \text{ s}^{-1}$) | $A_2$<br>(%) | $k_3$<br>($\times 10^{-3} \text{ s}^{-1}$) | $A_3$<br>(%) | $k_4$<br>($\times 10^{-3} \text{ s}^{-1}$) | $A_4$<br>(%) |
| --- | --- | --- | --- | --- | --- | --- | --- | --- |
| Fluorimeter | - | - | $87.6 \pm 18.8$ | $10 \pm 0.5$ | $15.0 \pm 0.8$ | $75 \pm 4.5$ | $2.66 \pm 0.06$ | $15 \pm 4$ |
| Stopped-flow | $535 \pm 70$ | $6 \pm 0.5$ | $65.2 \pm 7.3$ | $15 \pm 0.8$ | $9.1 \pm 0.2$ | $79 \pm 0.3$ | - | - |

Likewise, for the F21A mutant, the dissociation kinetics measured by stopped-flow were originally fitted to the sum of two exponential phases with a major slow phase (80%) of 25.8 ± 0.2 ×10^− 3^ s^-1^ and a minor fast phase (20%) of 423 ± 15 ×10^−3^ s^-1^. But when these data were fitted to the sum of three exponential phases, there was a major slow phase (66%) of 20.3 ± 0.2 ×10^−3^ s^-1^, a second faster phase (22%) of 91.8 ± 5.9 ×10^−3^ s^-1^, and third and fastest phase (12%) of 794 ± 68 ×10^−3^ s^-1^. This behaviour is similar to the dissociation kinetics observed for I19A by stopped-flow.

In contrast to these two fast-dissociating N-terminal mutants, the kinetics of the two fast-dissociating C-terminal mutants L41A and V44A was biphasic like the WT and there was no evidence of additional phases.

### The variable region of TCF7L2 is very sensitive to ionic strength

One of the major contributors to binding energy is electrostatic interactions, and the contribution can be investigated by varying the ionic strength of the buffer and measuring the changes in the association and dissociation rate constants. The magnitude of these variations can be assessed from the slopes of the plots of log(*k*_*on*_) and log(*k*_*off*_) versus log([NaCl]), referred to as Γ^on^_NaCl_ and Γ^off^_NaCl_ respectively (Chemes et al., 2011; Grucza et al., 2000; Vindigni et al., 1997). A negative value for Γ^on^_NaCl_ and a positive value for Γ^off^_NaCl_ indicates that the complex is destabilised by increasing ionic strength and therefore that electrostatic interactions are involved.

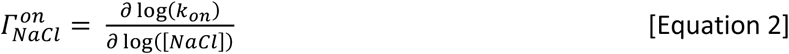

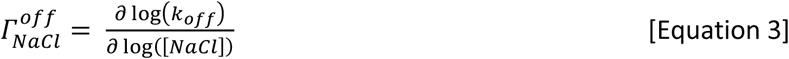

We used this method to test the interaction of β-catenin with both the WT TCF7L2 and the 5X variant, as we thought that the removal of five negatively charged residues might produce a significant change in any electrostatic interactions at the binding interface. We found for both WT and 5X that the association rate decreased with increasing ionic strength and the dissociation rate increased with increasing ionic strength (Fig. 7 and Table 3). However, the ionic strength dependence of the WT interaction was greater than that of the 5X mutant, and the 5X mutation had a much larger effect on the ionic strength dependence of dissociation (Γ^off,major^_NaCl_ and Γ^off,minor^_NaCl_) than on the salt dependence of association.

**Table 3.**
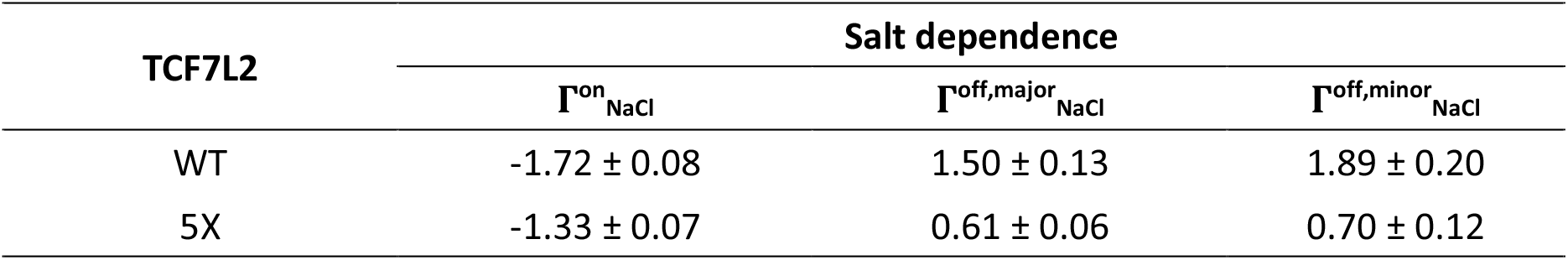
Ionic strength dependence of the interaction between β-catenin and TCF7L2 WT and 5X variant.

**Figure 7.**
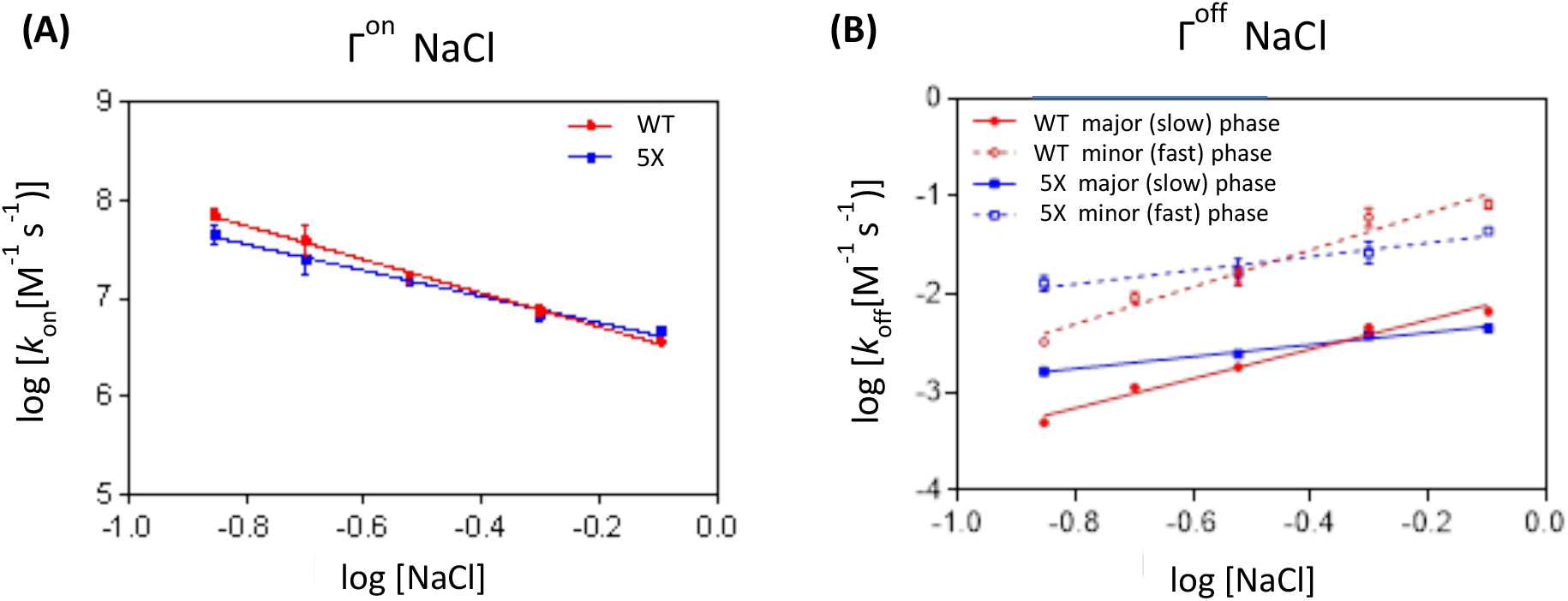
The effect of varying NaCl concentration on the kinetics of the interaction between β-catenin and the TCF7L2 WT and 5X variant. The effects of increasing NaCl concentration on the rate constant for association (A) and the rate constants for the two dissociation phases (B).

## Discussion

Our analysis of the kinetics of the β-catenin-TCF7L2 interaction was dissected by site-directed mutagenesis and uncovered a complex molecular recognition mechanism. We find that there are two interface key regions of TCF7L2 -an N-terminal binding region (residues 12 to 22) and a C-terminal binding region (residues 40 to 48) -that contribute significantly to the binding to β-catenin, and these are the regions that are the most similar between the different crystal structures. The single-exponential nature of the association kinetics makes it impossible to distinguish which section of this extended interface binds first, but it does suggest that any intermediates are of high energy indicating that that binding is highly co-operative. The trends observed in the alanine scan are consistent with an avidity mechanism of interaction between β-catenin and TCF7L2 (Olsen et al., 2017); the N- and C-terminal binding regions are connected by a flexible linker, each region binding its own cognate site on β-catenin, and once one site is bound the other region is held close in space to its corresponding site.

The central conformationally ‘variable’ region of TCF7L2 (residues 22-40), in which mutation had no effect on either the association or dissociation kinetics, can be regarded as a structurally malleable or “fuzzy” interaction generated by multiple rapidly exchanging high-entropy contacts eventually leading to a low entropy state (Fuxreiter, 2020; Toto et al., 2020). This type of fuzzy interaction, which lacks site-specific interactions, has also been seen recently for another binding partner of β-catenin, a 15-amino acid repeat region of APC (Rudeen et al., 2020). This fuzziness of binding of different partners involved in balancing the competing pathways of transcription and degradation may allow faster responses between the different pathways.

For the WT and alanine mutants in the C-terminal region of TCF7L2 the dissociation appeared to be biphasic. This behaviour can be explained as arising from sequential dissociation (Fig. 8A), whereby the C-terminal region always dissociates before the N-terminal region. In contrast, two alanine mutations in the N-terminal region of TC7L2, involving large truncations of hydrophobic sidechains (I19A and F21A), showed a four-phase dissociation reaction. It is likely that the interface-destabilising mutations in the N-terminal region cause a partial shift in flux through an alternative dissociation pathway, in which the N-terminal region dissociates before the C-terminal region, resulting in two additional kinetic phases being observed (Fig. 8B). This model is consistent with the observation that the alanine mutants in the C-terminal region in general have a greater effect on the dissociation rate constant than do the N-terminal alanine mutants.

**Figure 8.**
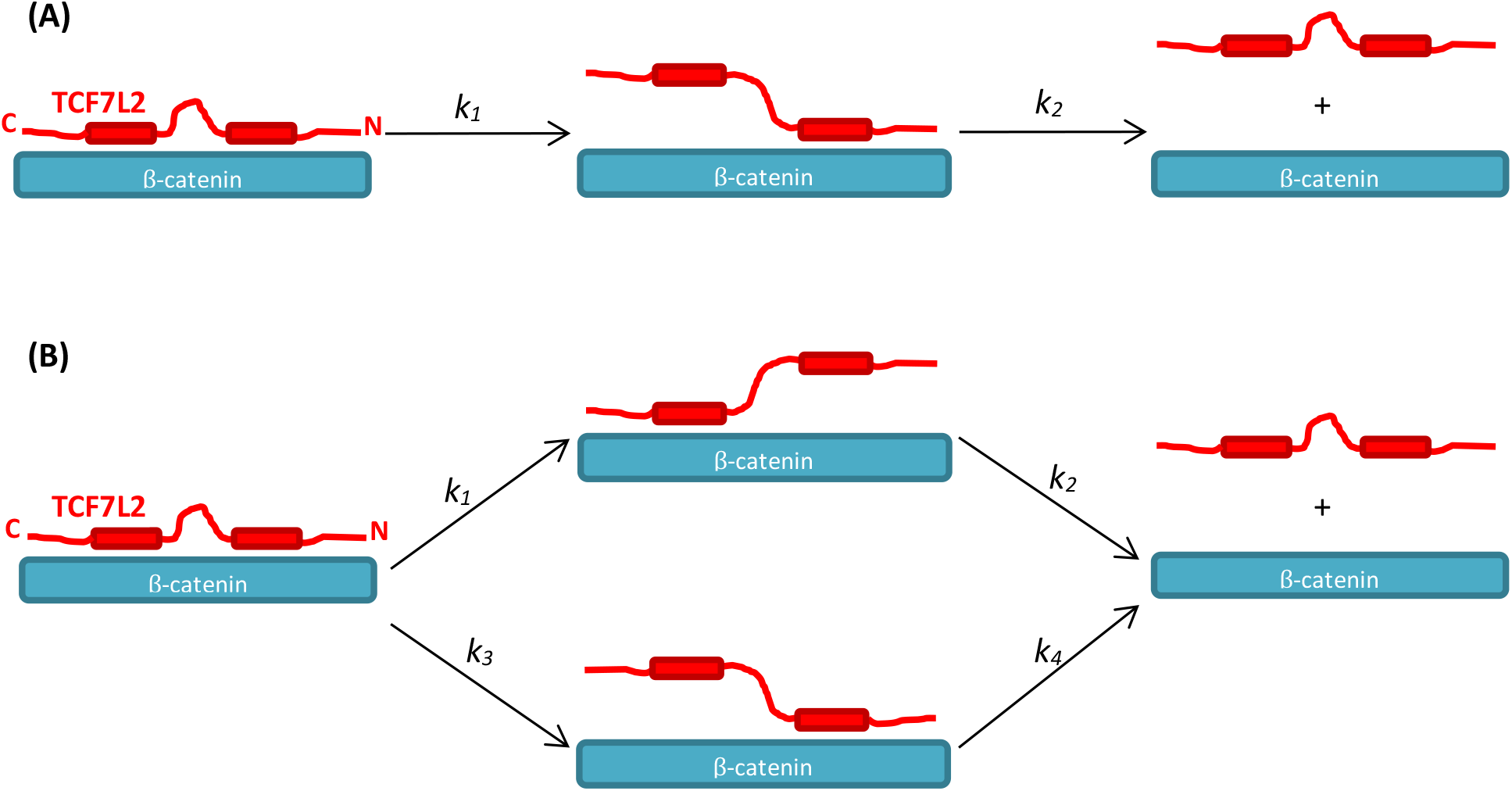
Schematic showing the proposed mechanism of dissociation of TCF7L2-β-catenin complex. TCF7L2 is in red and β-catenin in cyan. (A) Shows the sequential dissociation pathway in which the C-terminal region of TCF7L2 always dissociates before the N-terminal region. (B) For destabilising N-terminal mutants, there are two parallel dissociation pathways, in which the N-terminal region or the C-terminal region dissociates first.

The general trend for IDP interactions is that increasing ionic strength decreases *k*_*on*_ and has a minimal effect on *k*_*off*_ (Chemes et al., 2011; Dogan et al., 2014; Hemsath et al., 2005; Japrung et al., 2013; Rogers et al., 2013). In contrast, for TCF7L2-β-catenin there is a roughly equal effect of ionic strength on both dissociation and association: considering either of the two dissociation phases, Γ^off^_NaCl_ is similar in magnitude to Γ^on^_NaCl_. From the sum of Γ^on^_NaCl_ and Γ^off^_NaCl_ we can estimate a value of Γ^Kd,kin^_NaCl_ ≈ -3, which means that the TCF7L2-β-catenin complex is destabilised by three orders of magnitude for every order of magnitude increase in ionic strength. This sensitivity is relatively high compared to other IDP-protein interactions (e.g. Γ^Kd,kin^_NaCl_ ≈ -1 for the ACTR-NCBD interaction (Japrung et al., 2013); -1.4 > Γ^Kd,kin^_NaCl_ > -2.4 for various HPV E7 constructs binding to Rb (Chemes et al., 2011). There are parallels between our findings and those for the ACTR-NCBD interaction (Japrung et al., 2013). When ionic strength was increased, a reversal in the overall charge of NCBD occurred from positive to negative. The explanation given for this effect was that the highly positively charged NCBD attracts a shell of negatively charge chloride ions. At higher ionic strength, sufficient chloride ions are attracted to this shell to completely screen and eventually overwhelm the positive charge. We believe a similar behaviour occurs with TCF7L2-β-catenin, with a shell of positively charged sodium ions forming around the negatively charged variable region of TC7L2. The decrease in Γ^on^_NaCl_ upon mutation to alanine of all five charged residues can be readily explained, as increasing the ionic strength will have less of an effect on the interaction with β-catenin. The concept of an ionic shell/screen rationalises the unusually high Γ^off^_NaCl_ as well as the magnitude of the effect of the 5X Glu-to-Ala mutation on Γ^off^_NaCl_. As the variable region fluctuates between different conformations it periodically dissociates from the surface of β-catenin. While it is separated for β-catenin, the negatively charged residues can recruit a shell of positively charged sodium ions that inhibit re-association and promote complete dissociation. This effect increases in magnitude with increasing ionic strength and is ablated when the five charged residues are mutated to alanine in the 5X construct. Intriguingly the results of the recent single-molecule study of the E-cadherin-β-catenin interaction led to a similar suggestion, whereby there is continuous detachment and reattachment of local E-cadherin segments (Wiggers et al., 2021). We share the authors conclusion of a Velcro-like design of many weak contacts on top of a few persistent interactions.

The decrease in Γ^on^_NaCl_ upon mutation to alanine of all five charged residues can be readily explained, as increasing the ionic strength will have less of an effect on the interaction with β-catenin. The concept of an ionic shell/screen rationalises the unusually high Γ^off^_NaCl_ as well as the magnitude of the effect of the 5X mutation on Γ^off^_NaCl_. As the variable region fluctuates between different conformations it periodically dissociates from the surface of β-catenin. While it is separated for β-catenin, the negatively charged residues can recruit a shell of positively charged sodium ions that inhibit re-association and promote complete dissociation. This effect increases in magnitude with increasing ionic strength and is ablated when the five charged residues are mutated to alanine in the 5X construct.

The alternative pathways of binding and unbinding uncovered here for the ARM repeat protein β-catenin nicely mirror the alternative folding and unfolding pathways previously observed by our group and others for tandem-repeat proteins and previously predicted (Ferreiro et al., 2005, 2008; Hutton et al., 2015; Lowe and Itzhaki, 2007; Tsytlonok et al., 2013; Werbeck et al., 2008). These phenomena reflect the energy landscapes of the repeating architecture, in which the internal symmetry affords multiple paths of similar energies and, consequently, small perturbations such as conservative mutations are sufficient to shift the flux through each. Such behaviour contrasts with many globular proteins which often have rather complex topologies. In these proteins, there may only be one way to fold the protein, and therefore even drastic perturbations such as circular permutants can be insufficient to enable flux through any other routes (Lindberg and Oliveberg, 2007; Oliveberg and Wolynes, 2005; Otzen and Fersht, 1998). In the case of binding, the repeat protein-IDP interface is also strikingly different from most protein-protein interactions studied to date where there is no internal symmetry and consequently it is unlikely that there will be more than one for association/dissociation. In contrast, here we have looked at a relatively long, extended fragment of TCF7L2 binding to a very long interface comprising 11 repeat motifs. Consequently, the potential for the multiple sub-regions to ‘zip up’ sequentially and in different orders dependent on mutation is great. β-catenin is an important therapeutic target due to its deregulation in many diseases. However, both its long and relatively flat surface as well as the overlapping nature of its interfaces with favourable and unfavourable partner proteins have made it difficult to inhibit by conventional techniques. An understanding of the molecular recognition mechanisms involved, and in particular site-specific dissection of the complex pathways of dissociation will help us to direct drug design to more kinetically accessible interfaces of β-catenin and thereby most effectively disrupt or displace binding partners partner proteins or displace or displace binding partners and inhibit its transcriptional activity.

