## Supplemental Information for "Parallel and sequential pathways of molecular recognition of a tandem-repeat protein and its intrinsically disordered binding partner"

### Supplementary Information

**Table S1: Dissociation constants of TCF7L2 variants binding to β-catenin**

| Construct | Method | $K_d$ (nM) | Stoichiometry |
|---|---|---|---|
| TCF7L2 (1-53) (Omer *et. al.*) | ELISA | 15 ± 6 | n.d. |
| TCF7L2 (1-57) (Sun & Weis) | ITC | 16 ± 3 | 2 |
| TCF7L2 (1-54) | ITC | 13 | 1 |
| WT S31C | ITC | 15 | 1 |
| WT fluorescently labelled | ITC | 28 | 1 |

Data from (Sun and Weis, 2011) are for TCF7L2 (1-57) binding to full-length β-catenin. Experiments were performed in 25 mM Tris-HCl pH 8.9, 100 mM NaCl, 2 mM DTT at 30°C, whereas the experiments of Sun and Weis were performed in 25 mM Tris-HCl pH 8.8, 100 mM NaCl, 2 mM DTT at 30°C.

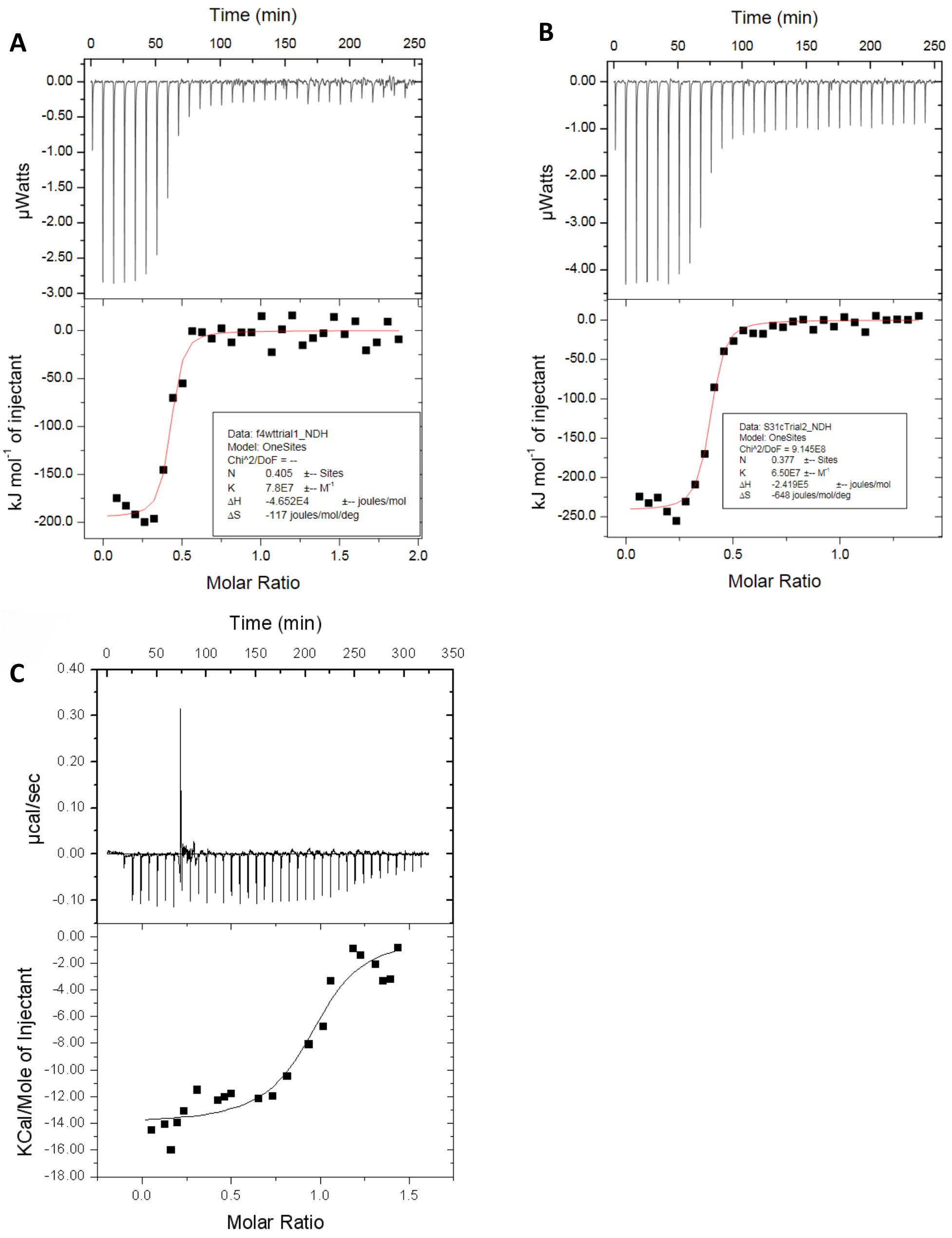


**Figure S1. ITC analysis of TCF7L2 constructs binding to β-catenin.** The top panels show the heat signal obtained from a series of injections of different TCF7L2 into the ITC cell containing β-catenin, and the bottom panels show the binding curves calculated using the One-Site fitting model using the Origin software package. (A), (B) and (C) are TCF7L2 (1-54), "WT" and the fluorescein-labelled WT construct binding to β-catenin, respectively. Experiments were performed in 25 mM Tris-HCl pH 8.9, 100 mM NaCl, 2 mM DTT at 30°C.
